# Changes in mangrove tree mortality, forest canopy, and aboveground biomass accumulation rates following the 2017 hurricane season in Puerto Rico and the role of urbanization

**DOI:** 10.1101/425140

**Authors:** Benjamin L. Branoff

**Affiliations:** Department of Biology University of Puerto Rico-Río Piedras San Juan, Puerto Rico, 00931, U.S.A

## Abstract

Mangrove ecosystem responses to tropical cyclones have been well documented over the last half a century, resulting in repeated measures of tree mortality, aboveground biomass reduction, and recovery by species, size, and geomorphology. However, no studies have investigated the role of urbanization in mangrove hurricane resistance and resilience, despite increasing urbanization of tropical shorelines. This study gauges the initial response and short-term recovery of Puerto Rico’s mangroves along well defined and quantified urban gradients following the 2017 hurricane season. Survival probability of tagged trees decreased with time, and the mean mortality across all sites was 22% after eleven months. Mean canopy closure loss was 51% one month after the hurricanes, and closure rates also decreased with time following the storms. Aboveground biomass accumulation decreased by 3.5 kg yr^-1^ per tree, corresponding to a reduction of 4.5 Mg ha^-1^ yr^-1^ at the stand level. One year later, the mangroves have recovered to 72% canopy closure and to nearly 60% of their pre-storm growth rates. No connection to urbanization could be detected in the measured dynamics. Instead, species, size and geomorphology were found to play a role. Larger trees suffered 25% more mortality than smaller size classes, and *Laguncularia racemosa* suffered 11% less mortality than other species. Hydro-geomorphology was also found to play a role, with forests in tidally restricted canals experiencing more canopy loss but faster recovery than open embayment systems. These findings suggest size, species, and geomorphology are important in mangrove resistance and resilience to tropical storms, and that urbanization does not play a role. Managing mangrove ecosystems for optimal shoreline protection will depend upon knowing which forests are at greatest risk in a future of increasing urbanization.

## Introduction

Tropical cyclones are sources of repetitive disturbance in coastal communities around the world, with US$26 billion spent annually on damages to property and infrastructure inflicted by these storms (Mendelsohn et al. 2012). This figure is expected to double by 2100 due to an ongoing migration of the global population towards tropical cities, putting more lives and property within the reach of cyclone disturbance (Mendelsohn et al. 2012). Coastal wetlands have been shown to reduce the damages to infrastructure and property caused by tropical cyclones (Costanza et al. 2008), and mangrove forests are singled out as especially effective in coastal protection (Das and Vincent 2009; Narayan et al. 2011; Marois and Mitsch 2015). But mangroves in urban landscapes, where their service as coastal protection is most valuable, are diminishing faster than the global average (Branoff 2017). Further, although multiple studies have shown how mangroves respond to tropical storm events (Wadsworth 1959; Smith et al. 1994; A. H. Baldwin et al. 1995; McCoy et al. 1996; Smith III et al. 2009; Daniel Imbert 2018), none have evaluated how urbanization influences this response. Thus, the management of urban mangroves towards optimal provisioning of protective services cannot effectively evaluate the role of the urban landscape.

Overall, initial mangrove mortality following tropical storms has ranged from 25%-90%, with the variation being attributed to differences in species, tree size, and hydro-geomorphology, in addition to storm intensity and location (Craighead and Gilbert 1962; Roth 1992; Smith et al. 1994; Armentano et al. 1995; Sherman et al. 2001). Where size was accounted for, studies have almost always shown larger trees to be most susceptible to both partial and complete mortality (Roth 1992; Smith et al. 1994; Doyle et al. 1995; McCoy et al. 1996), with the exception of one study showing no relationship (Sherman et al. 2001). For variations within species, conclusions are more often conflicting than in agreement. Following hurricane Andrew in Florida, *R. mangle (Rhizophora mangle)* was found to suffer the highest mortality, and *L. racemosa (Laguncularia racemosa)* the least (A. H. Baldwin et al. 1995), but other studies of the same hurricane in Florida found the opposite (Doyle et al. 1995; McCoy et al. 1996). Still another study of the same storm in the Dominican Republic found *L. racemosa* to be the least affected, and *A. germinans (Avicennia germinans)* the most (Sherman et al. 2001). Other studies for other storms in other locations have found variations in species susceptibilities (Wadsworth 1959; Smith et al. 1994; Daniel Imbert 2018). This conflict might be explained by differences in habitat and hydro-geomorphology, both of which have also been found to play a role in storm related tree mortality and forest recovery (Sherman et al. 2001; Smith III et al. 2009; Daniel Imbert 2018).

Recovery patterns show trends within the same predictors of species, size, and geomorphology, again with conflicting conclusions (Roth 1992; A. H. Baldwin et al. 1995; Sherman et al. 2001; Daniel Imbert 2018). One long-term study of post-hurricane Caribbean mangroves suggest a recovery time to pre-storm similarity of 10-25 years, if at all (Daniel Imbert 2018), and modelling approaches generally agree (Lugo et al. 1976; Doyle and Girod 1997). However, the influence of these disturbances on mangroves is so ubiquitous, that It has been hypothesized they permanently restrict the height of Caribbean mangroves (Odum and Pigeon 1970; Lugo and Snedaker 1974). Further, due to the above stated differences in susceptibility and recovery, these forests are thought to be constantly shifting composition in response to periodic tropical storms (Smith et al. 1994; A. Baldwin et al. 2001; Piou et al. 2006). Thus, depending upon storm intensity, forest structure, and geomorphology, it is possible to provide limited predictions on the potential effects a storm will have on mangrove forests, as well as recovery pathways. But mangroves increasingly inhabit mixed-use landscapes (Thomas et al. 2017), and urbanization has been absent from consideration in any of the previous studies on Caribbean mangrove hurricane response.

Mangroves have been shown to exhibit greater mortality than other forests following storms (Armentano et al. 1995), so urban mangroves may be especially susceptible to tropical storm disturbance. Further, the Caribbean has been highlighted as a biodiversity hot-spot predicted to see a larger than average urbanization rate by 2030 (Seto et al. 2012). If this forecast is accurate, and if urban mangroves are less resistant and resilient than other forests, it could lead to diminished protective services of urban mangroves, and thus more susceptible human communities along tropical urban coastlines.

This study aims to capture the response of Puerto Rico’s urban mangrove forests following two separate tropical cyclone events in 2017. Hurricane Irma was the strongest hurricane ever in the open Atlantic Ocean, passing within 93 kilometers of Puerto Rico’s north coast on September 6^th^ with maximum wind speeds on the island of 110 km/h (Cangialosi et al. 2018) (Figure 1a). Two weeks later, on September 20^th^, Hurricane Maria made landfall along the southeastern coast of Puerto Rico with maximum winds of 250 km/h (Pasch et al. 2018). The storm’s center tracked northwest across the island for eight hours, leaving with maximum winds of 175 km/h. Damage to infrastructure and property from hurricane Maria was estimated at US$65 - US$115 billion (Pasch et al. 2018).

**Figure 1.**
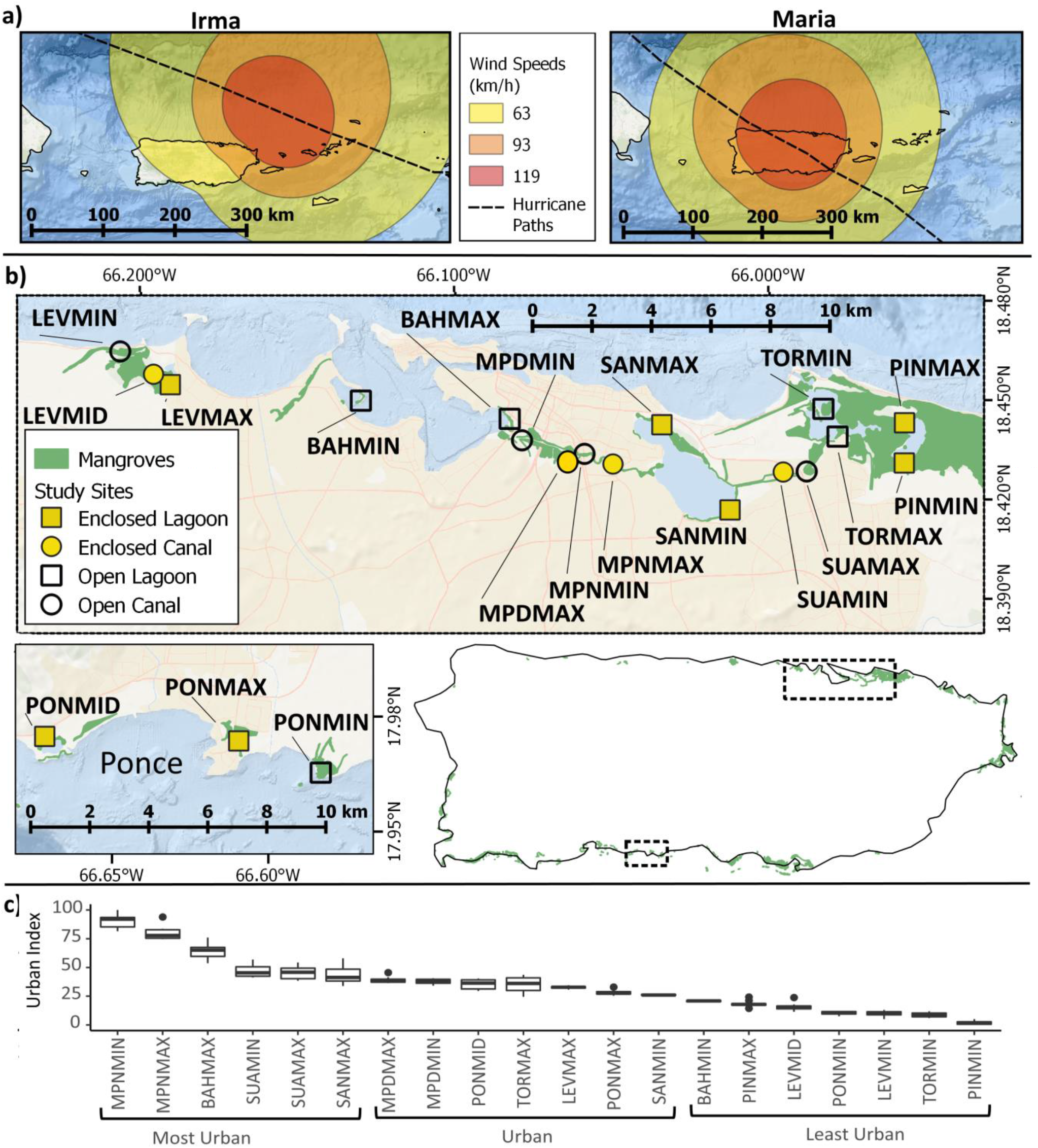
Hurricanes Irma and Maria subjected Puerto Rico to hurricane force winds within two weeks of each other (a), with Maria being the worst natural disaster in Puerto Rico’s history. Study sites consisted of 20 one-hectare forested mangrove sites in three watersheds of Puerto Rico, representing one of four potential hydro-geomorphologies (b). Maria passed within 25 km of Levittown sites, and with 45 km of Ponce sites. Sites were placed along a gradient of urbanization, as defined by an urban index, to maximize the difference between the most urban and least urban sites (c).

This study uses previously tagged trees and repetitive dendrometer and canopy closure measurements to test for differences in initial mortality and canopy loss, as well as short-term recovery across species, size, hydro-geomorphologies, urbanization, and storm wind power in Puerto Rico’s mangroves. Results will be used to gauge the predicted recovery times in mangroves across the island, as well as propose potential management considerations for optimizing the provisioning of protective services to the island’s infrastructure and inhabitants.

## Methods

Study sites consisted of 20 one-hectare forested mangrove areas along quantified urban gradients in three watersheds of Puerto Rico (Figure 1b, Table 1). Urbanization at each site was defined by an urban index (Branoff 2018)(Figure 1c), which was calculated using surrounding (within 0.5 km) population density, road length, and urban, open water, vegetated, and mangrove land covers. The most urban sites were classified as those in the 75^th^ percentile of the urban index, the least urban were those in the 25^th^ percentile, and urban were all other sites within the 25^th^ to 75^th^ percentiles of the urban index. All sites were located along a shoreline, thus restricting their classification as fringe systems, but hydro-geomorphological settings were classified as partially restricted or fully open to tidal influence, and as canal or embayment (e.g. lagoon, bay, ocean etc.) as described by (Branoff 2018). Ten 5 m radius circular plots were established at each site and their vegetation structural and compositional characteristics are described in (Branoff and Martinuzzi 2018). In general, *L. racemosa* represents 51% of the trees in these forests, followed by *R. mangle* at 29%, *A. germinans* at 9%, and *Thespesia populnea* at 7.5%. The remaining trees are represented by twenty-five additional, non-halophyte species. There were no differences in species composition, dbh, stem density, basal area, or aboveground biomass between watersheds. Field measurements of tree size and canopy closure as described below commenced on different dates but were taken concurrently thereafter, with an average frequency of 100 days.

**Table 1.** Site abbreviations used throughout the study and their corresponding locations

| Watershed | Site | Description | Latitude | Longitude |
| --- | --- | --- | --- | --- |
| San Juan Bay Estuary - Río Bayamón - Río Hondo to the Río<br>Puerto Nuevo - Río Piedras | BAHMAX | San Juan Bay at Parque Central | 18.44434 | -66.08269 |
|  | BAHMIN | San Juan Bay at Peninsula la Esperanza | 18.44983 | -66.13007 |
|  | MPDMAX | Dredged portion of Martín Peña Canal, near Hato Rey ferry terminal | 18.43328 | -66.06349 |
|  | MPDMIN | Dredged portion of Martín Peña Canal, junction with Río Piedras | 18.43786 | -66.07877 |
|  | MPNMAX | Undredged portion of Martín Peña canal at calle Pepe Díaz | 18.43070 | -66.04982 |
|  | MPNMIN | Undredged portion of Martín Peña canal at PR highway 1 | 18.43376 | -66.05888 |
|  | PINMAX | Piñones lagoon at entrance to Bosque Estatal de Piñones | 18.44315 | -65.95701 |
|  | PINMIN | Along southern shore of Piñones lagoon | 18.43101 | -65.95708 |
|  | SANMAX | San José lagoon at calle mar amarillo | 18.44256 | -66.03424 |
|  | SANMIN | San José lagoon at southern bank of San Antón creek | 18.41683 | -66.01256 |
|  | SUAMAX | Southern bank of Suarez canal, just east of PR hwy 26 bridge | 18.42845 | -65.98815 |
|  | SUAMIN | Northern bank of Suarez canal, 0.5 km west of PR hwy 26 bridge | 18.42828 | -65.99562 |
|  | TORMAX | Torrecillas lagoon, just north of calle Sevilla | 18.43898 | -65.97833 |
|  | TORMIN | Torrecillas lagoon at Punta Larga island | 18.44736 | -65.98283 |
| Levittown<br>– Río la<br>Plata | LEVMAX | Western shore of Levittown lakes | 18.45467 | -66.19073 |
|  | LEVMID | Southern bank of Levittown lakes drainage creek | 18.45780 | -66.19606 |
|  | LEVMIN | Northern bank of Río Cocal, 1 km southwest of its mouth | 18.46470 | -66.20667 |
| Ponce - Río<br>Inabón to<br>the Río Loco | PONMAX | Northeast corner of intersection of PR hwy 12 and PR hwy 123 in Ponce | 17.97259 | -66.60947 |
|  | PONMID | Northern shore of laguna de Salinas at the cuchara nature preserve, Ponce | 17.97397 | -66.67151 |
|  | PONMIN | Punta Cabullones, Ponce, eastern shore of inlet | 17.96294 | -66.58324 |

Tree growth was measured using stainless steel band dendrometers as described by Cattelino et al (1986). Ten dendrometers were installed at each site from May to July of 2017, resulting in 200 dendrometers across the three watersheds. Trees were selected to represent as many species and sizes of each species at each site as possible. The minimum, median, mean, and maximum diameter at breast height (dbh) of dendrometer trees were 3, 13, 15.2, and 54 cm, respectively. The same statistics for the 9,400 trees measured at all sites were 1, 4.6, 6.7, and 54 cm. Thus, the dendrometers represent a bias towards larger trees due to the difficulty of accurately banding those smaller than 5 cm diameter. Tree diameters were classified into size classes of four equal quantiles representing 25% each of the total distribution. Collar starting positions were marked on the band by scratching with a sharp knife. Incremental growth was measured using a caliper as the distance between the starting scratch and the position of the collar. Trees were determined dead if they exhibited shedding bark, no leaves, and if scratching did not produce green cambium tissue. Dendrometers were removed from dead trees and placed on the nearest similar tree in the plot. If the same size class of the same species could not be found in that plot, it was placed on one in another plot. If that could not be found, it was placed on the same size class of another species in the same plot.

Tree mortality was tabulated as alive or dead with each visit and the length in days since hurricane Maria was calculated for each confirmed death. The resulting time-series of deaths was used to create Kaplan-Meier survival curves (Kaplan and Meier 1958; Swinscow and Campbell 2002) for each grouping of trees using the *survfit* function from the survival package (Therneau and Grambsch 2000). This function calculates the non-parametric probability of a patient, in this case a tree, surviving past a certain event, in this case hurricane Maria, based upon the time of death for similar trees. Differences in survival curves among groupings were inferred from log-rank tests (Harrington and Fleming 1982) as calculated from the *survdiff* function of the same package.

Growth in diameter was converted to aboveground biomass accumulation using allometric equations specific to each species and size class. For the three true mangrove species of *A. germinans*, *L. racemosa*, and *R. mangle*, equations were derived from three sources on Caribbean mangroves, and the mean was used when equations overlapped (D Imbert 1989; Fromard et al. 1998; Smith and Whelan 2006). When no value was available for greater size classes, a general equation for mangrove habitats was used from Chave et al. (2005). This equation was also used for non-mangrove species in combination with specific gravities derived from Reyes et al. (1992). Growth rates were taken as the difference in measurement values from one date to the next over the length in days between measurements. This was then converted to a unit of kg/yr by multiplying by 365. Because all sites could not be measured at once or during every measurement campaign, measurements were interpolated to a monthly frequency based on calculated rates. Thus, the measurement for an interpolated date was taken as the calculated rate for that period multiplied by the time length since the previous measurement. Aboveground biomass accumulation for the entire period after the storm was calculated by integrating the area under a loess curve fit over the monthly interpolated growth rates (Odum and Odum 2000). Aboveground biomass accumulation before the storm could not be integrated due to too few measurements and was instead taken as the mean growth rate multiplied by the time duration. Aboveground biomass accumulation at the stand level was calculated for each site by taking the calculated accumulation rates for each species in each size class at each site, and multiplying by the number of trees of each species in each size class per hectare at each site, as taken from Branoff and Martinuzzi (2018). This resulted in stand level aboveground biomass accumulation in units of Mg ha^-1^ yr^-1^.

Canopy closure before and after the hurricane was assessed using two different methodologies, LiDAR and hemispherical photos, respectively. Closure before the hurricane was assessed through LiDAR data obtained in March of 2017 for the San Juan sites as part of a NASA GLiHT campaign (Cook et al. 2013; Branoff and Martinuzzi 2018). LiDAR data has previously been shown to slightly overestimate measurements from hemispherical photos, with a mean error of 4-7% (Korhonen et al. 2011). Closure from LiDAR data was taken as the fractional coverage of trees, or the percentage of first returns sensed as trees. Canopy closure following the hurricanes was measured using semi-hemispherical photos taken from the ground (Evans and Coombe 1959; Valverde and Silvertown 1997). Photography began in October of 2017 and terminated in August of 2018. Photos were taken using a GoPro Hero camera with a 170° field of view. The camera was placed at the center of each plot at a height of 50 cm and oriented so that the bottom of the lens pointed north. Photos were taken just after dawn, before dusk, or during overcast conditions, when possible, to avoid interference from direct sunlight.

Photos were processed as follows in the R programming language (Yan et al. 2011) to produce binary images of closed and open canopy. The blue channel of each photo was used to reduce variance (Brusa and Bunker 2014), and was separated using the *channel* function from the EBImage package (Pau et al. 2010). The Otsu threshold is that which optimally creates a binary image from a greyscale image (Otsu 1979). In this case, the threshold seeks to automatically detect which pixels are canopy, and which are sky, depending upon their level of grey. This was done for each canopy photo using the *otsu* function also from EBImage. Each binary image was then visually inspected to ensure proper representation of the original. When errors in thresholding were detected, thresholds were incrementally increased or decreased, depending upon the error, until proper representation was achieved. If errors persisted, the problematic regions were manually adjusted to either black or white in the imageJ software (Schneider et al. 2012). Canopy closure was then calculated as the percentage of canopy pixels in each binary image, or the number of pixels with a value of one, over the total number of pixels. As with tree growth, canopy closure was interpolated to a monthly frequency based on the rate of change between measurements.

Distance from each study site to the closest passing of hurricane Maria’s center was calculated using the *gDistance* function from the rgeos package (Bivand and Rundel 2017) and a shapefile of the storm’s track from the national hurricane center (National Hurricane Center 2017). Wind power in units of hMJ m^-3^ at each site were taken from Figure 2a of Van Beusekom et al. (2018), which represents the total gale wind kinetic energy from both hurricanes, taking into account topography and estimated wind speeds. Wind power was extracted from this dataset using the *extract* function from the raster package (Hijmans 2016).

**Figure 2.**
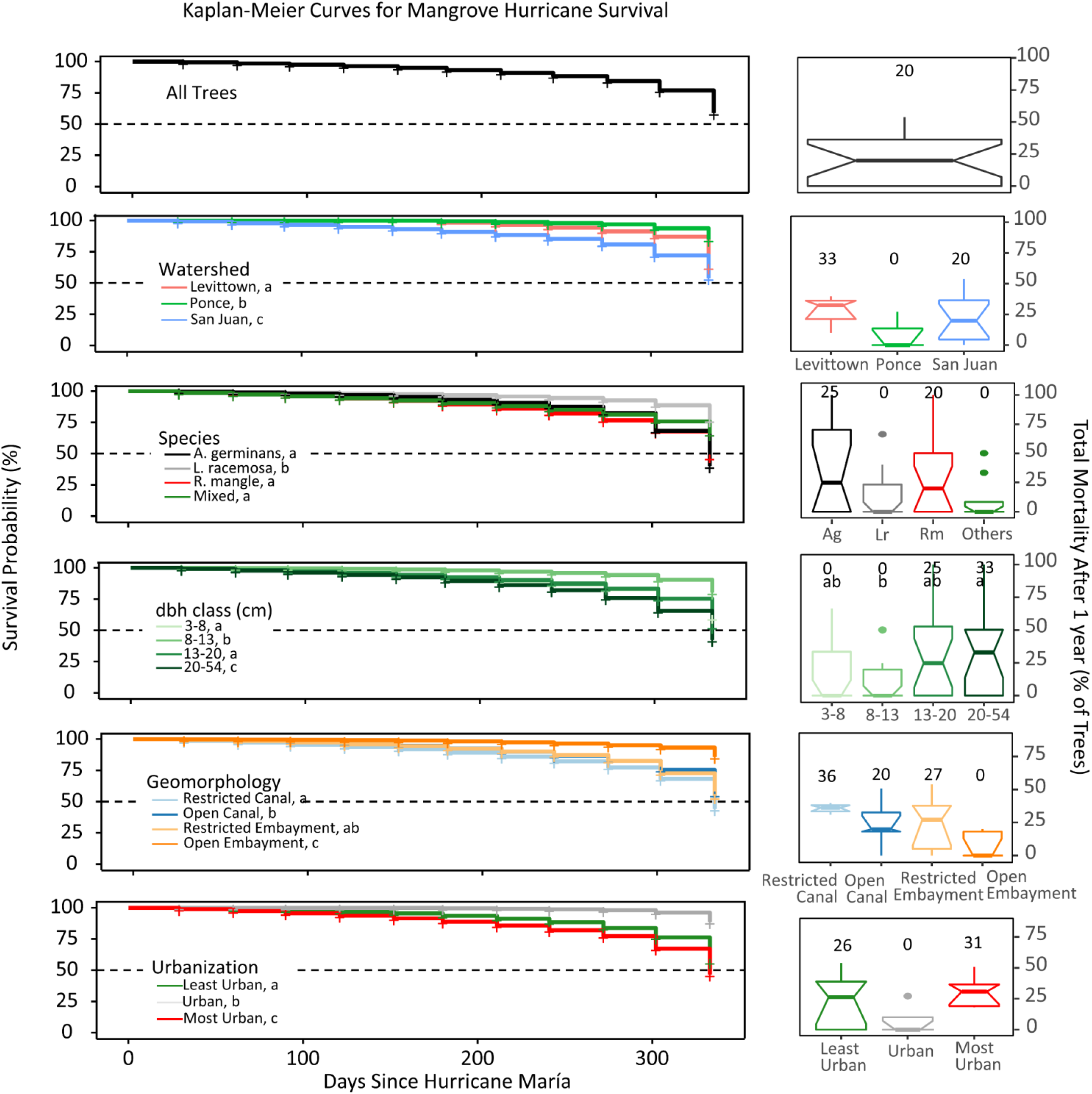
Kaplan-Meier survival probability curves with time since hurricane Maria for each group of trees (left), and the final mortality percentage of all groups after one year (right). Statistically similar groups are denoted by the same letter when differences were present, and median values for each group are shown over boxplots. Survival probability remained above 90% until 250 days following the storm, when it began to fall precipitously to a value of 60% at 315 days. *L. racemosa* trees maintained a higher survival probability, as did medium sized trees, those in open embayments, of intermediate urbanization, and those in Ponce. Mortality after 11 months was highest in the largest trees and lowest in *L. racemosa* and non-mangrove species.

All data analysis was done in R. Initial canopy loss, mortality, and growth rates were compared between species, size classes, urbanization, and geomorphology using analysis of variance (ANOVA) and subsequent post-hoc Tukey honest significant differences through the *aov* and *TukeyHSD* functions in base R. Data were plotted through the *ggplot* function from the ggplot2 package (Wickham 2009), and linear and logarithmic models were constructed through the *lm* function, also from base R.

## Results

### Mortality

Survival probability remained above 90% for the first 250 days following hurricane Maria but dropped sharply to 60% by day 315 (Figure 2). Survival curves were different among all groupings (log-rank test; p < 0.05), with intermediately urban *L. racemosa* trees of small size in open embayments of Ponce expressing the highest overall survival probability over the course of the year. As of eleven months following the hurricanes, overall mean mortality across all tagged trees was 22% and the only significant differences found were between size classes (ANOVA; p< 0.05) (Figure 2, Table 2). The largest size class of 20-54 cm dbh experienced the greatest mean mortality rate of 33%, which was significantly different than the intermediate class of 8-13 cm at 9.2% (ANOVA; difference = 24%, p < 0.05). The 13-20 cm class experienced a mean mortality of 32%, followed by the smallest class of 3-8 cm at 17%. Differences by species were barely insignificant (ANOVA; p = 0.07). *A. germinans* exhibited the highest mean mortality of 38%, followed by *R. mangle* at 28%, *L. racemosa* at 14%, and non-mangrove species at 11%. Mortality was also insignificant by watershed, urbanization (ANOVA; p > 0.5), distance from storm track (linear model; p > 0.1), or total wind energy (linear model; p>0.5). Still, San Juan and Levittown each experienced more than double mean mortality, at 28% and 23%, respectively, in comparison with Ponce at 9%.

**Table 2.**
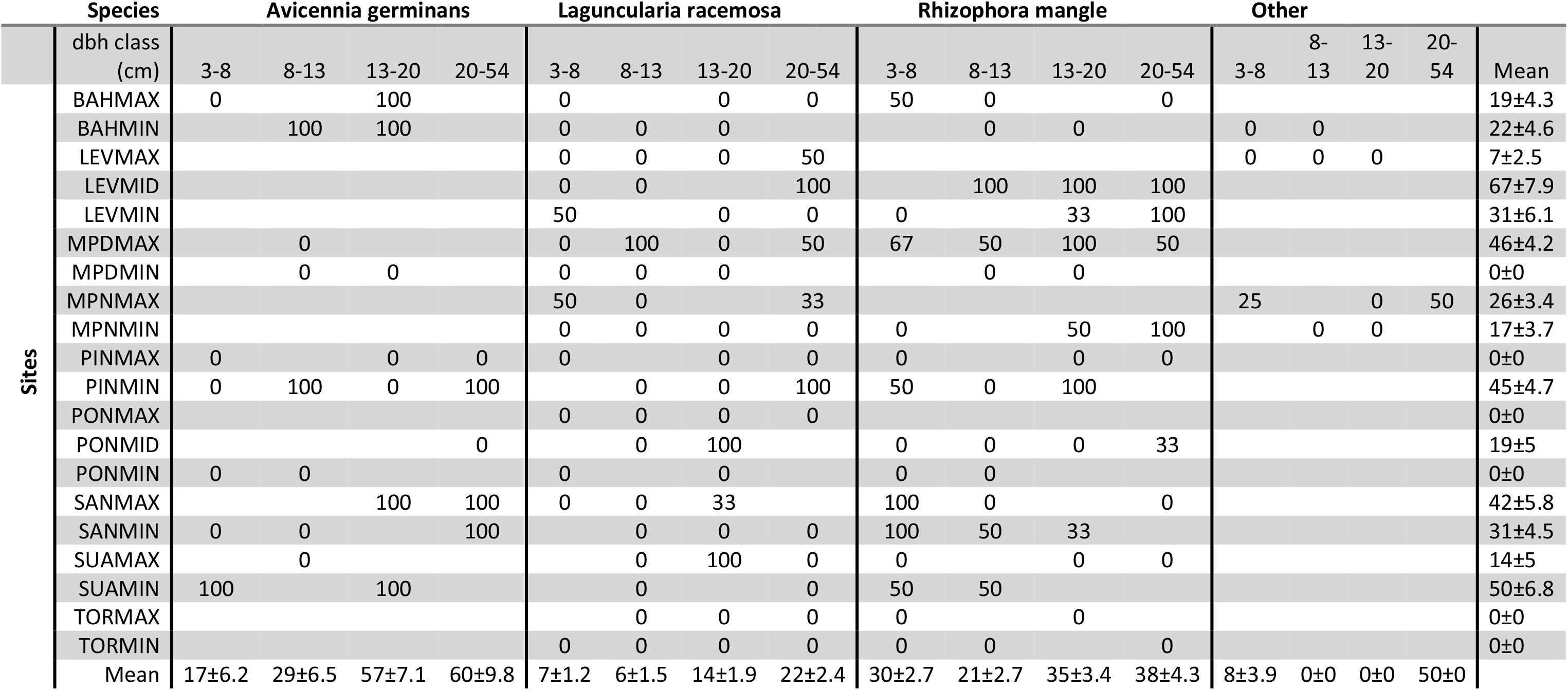
Mean percent tree mortality after eleven months from ten trees at each site, grouped by species and size. Means and standard errors for each size class and each site are given on the bottom and right, respectively. Site locations are demonstrated in Figure 1 and Table 1.

### Canopy Closure

Mean canopy closure loss one months after hurricane Maria could only be attained at San Juan sites due to LiDAR availability and was 51% (Figure 3). There were no differences in mean canopy loss between average tree size (ANOVA; p > 0.1) or urbanization (ANOVA; p > 0.5). But sites dominated by *A. germinans* lost 11% more canopy closure than sites dominated by *L. racemosa* (ANOVA; p < 0.01). Further, canopy loss increased linearly with percent of stand biomass as *A. germinans*, at a rate of 0.2% canopy loss for every percent of stand biomass as *A. germinans* (linear model; p < 0.001). Also, forests in tidally open systems lost 10% less canopy closure than those in restricted systems (ANOVA; p < 0.001). As with mortality, there was also no relationship between distance to storm track or cumulative wind energy with canopy loss following the storm (linear models; p > 0.5).Overall canopy recovery averaged 2% per month, but this rate decreased progressively with time following the hurricane (Figure 3). Recovery was fastest for the first three months following the hurricane, at 3.4% closure per month. From the fourth to the sixth month, recovery was 2.8% per month, from the sixth to the ninth month it was 1.5% per month, and from the ninth to the eleventh month it was 1.3% per month. Overall canopy closure one year after the storms was 72% (Table 3). There were no differences in overall canopy recovery rates by geomorphology or watershed (ANOVA; p > 0.5), and as with initial canopy loss, there also were no differences between mean tree sizes or urbanization (ANOVA; p > 0.5). There were however, differences in recovery rates by species composition, with forests dominated by *R. mangle* expressing the slowest recovery rate at 1.1%, slower than all other forest types (ANOVA; mean difference = 1%, p < 0.05). *L. racemosa* showed the highest recovery rate at 2.4% per month, followed by mixed forests and forests of *A. germinans*, both at 2.0%. Thus, *L. racemosa* dominated forests are forecasted to recover fastest to pre-hurricane canopy closure, reaching 80%, 85%, and 95% closure within 1.7, 4, and 6 years, respectively. All other forests will likely take longer than twenty years to reach these milestones.

**Figure 3.**
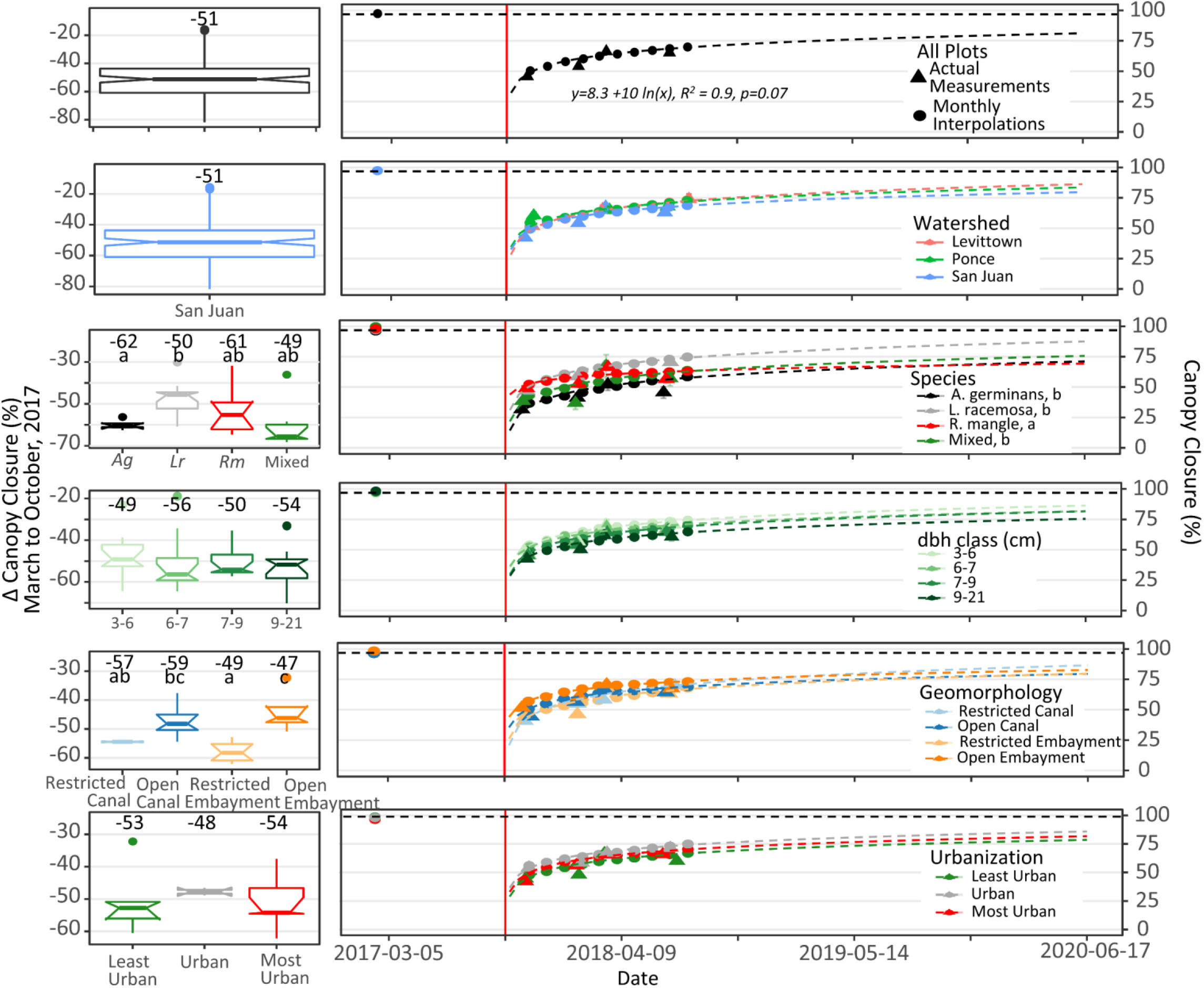
The change in canopy closure one-month after hurricane Maria (left), and canopy closure with time (right), grouped by watershed, species, diameter, geomorphology, and urbanization. The vertical red line is hurricane Maria. Triangles ns of actual measurements and circles represent interpolated monthly values. Statistically similar groups are denoted by the same letter when differences were present, and median values for each group are shown over boxplots. Measurements before the storm were obtained by LiDAR, which was only available for San Juan, and those after by hemispherical photos. Canopy closure loss was highest in *A. germinans* forests and in those of tidally restricted geomorphologies. The only difference in canopy closure recovery rates was detected in forests dominated by *A. germinans R. mangle*, which closed slower than all other forest types. Overall, closure to 80, 90, and 95 % can be expected in 3.6, 9.7, and 16 years, respectively, but some forests may take considerably longer.

**Table 3.** Mean and standard error percent canopy closure in the forests as measured by LiDAR before the hurricanes, and as measured by semi-hemispherical photos at ten days and eleven months following the hurricanes.

| Site | Before<br>Maria | Ten<br>Days<br>After<br>Maria | Eleven<br>Months<br>After<br>Maria |
| --- | --- | --- | --- |
| BAHMAX | 97±0.8 | 51±4.9 | 74±4.5 |
| BAHMIN |  | 57±5.5 | 59±7.6 |
| LEVMAX |  | 69±6.8 | 93±0.9 |
| LEVMID |  | 41±3.2 | 70±7.7 |
| LEVMIN |  | 37±2.4 | 69±10.3 |
| MPDMAX | 96±0.9 | 42±3.1 | 59±9.2 |
| MPDMIN | 98±0.6 | 49±4.5 | 70±5.7 |
| MPNMAX | 97±0.9 | 43±3.9 | 75±8.3 |
| MPNMIN | 94±2 | 57±4.1 | 77±5.6 |
| PINMAX | 98±0.6 | 45±4.2 | 82±6.7 |
| PINMIN | 95±1.2 | 35±3.4 | 55±7.8 |
| PONMAX |  | 41±8.5 | 87±2.1 |
| PONMID |  | 57±3.4 | 61±6.7 |
| PONMIN |  | 60±4.7 | 83±5.7 |
| SANMAX | 98±0.5 | 35±3.9 | 59±9 |
| SANMIN | 98±0.3 | 42±2.7 | 63±7.1 |
| SUAMAX | 95±1.4 | 48±4.5 | 78±5.8 |
| SUAMIN | 95±0.9 | 40±3.9 | 79±5.4 |
| TORMAX | 97±1 | 50±2.9 | 81±4.8 |
| TORMIN | 99±0.3 | 48±1.9 | 74±3.4 |
| Mean | 97±0.4 | 47±2 | 72±2.3 |

### Growth

Mean and median tree growth rates dropped by 2.3 kg/yr and 0.6 kg/yr, respectively, from the first two measurement made before the hurricanes to the first measurement after (Figure 4). Differences in mean change in growth rates from before and one month after the storms were detected between species, size, and geomorphology. Non-mangrove species accelerated growth following the storms, resulting in a significant difference in the immediate (within one month of hurricane Maria) change of growth rates between them and all mangrove species (ANOVA; mean difference = 14 kg yr^-1^, p < 0.05). There was also a difference in sizes, with the two largest size classes slowing growth and the two smallest classes accelerating (ANOVA; mean difference = 8.8 kg yr^-1^, p < 0.05). Likewise, tidally restricted canal trees also accelerated growth following the storm, while all other geomorphologies slowed, resulting in a significant difference between restricted canals and open embayments (ANOVA; difference = 7.3 kg yr^-1^, p < 0.05).

**Figure 4.**
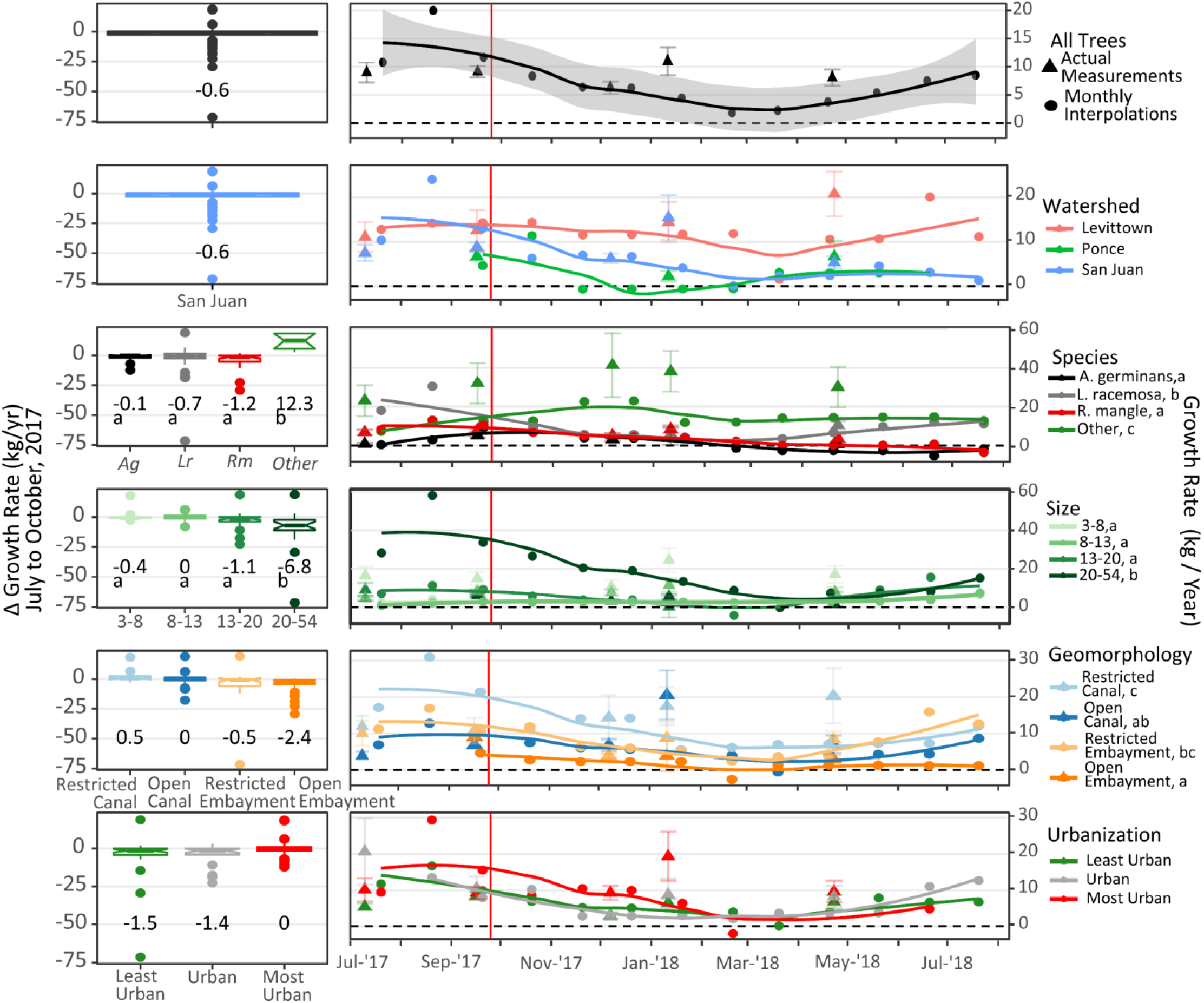
The change in individual tree growth rates from before and one-month after hurricane Maria (left), and growth with time (right) as grouped by watershed, species, diameter, geomorphology, and urbanization. Triangles represent means of actual measurements at the midpoint between measurement dates. Circles represent interpolated monthly values. The shaded area and the error bars represent standard error of the mean. The vertical red line is the date of hurricane Maria. Medians are shown above boxplots and letters designate statistically similar groups when differences were present. In the case of growth with time, letters designate similar growth rates after hurricane Maria. After the hurricane, non-mangrove trees grew faster than mangroves, and although the largest trees saw the steepest reduction in growth rates, they still grew faster than the smaller trees. Also, trees in restricted hydro-geomorphologies grew faster than those in open systems.

In comparing the year of growth following the storm, there was an overall decrease in aboveground biomass accumulation compared to before the storm. Integration of growth rates before the storm was not done for lack of measurements, but the mean of growth rate measurements before the storm was 9 kg/yr. Integration of rates after the storm resulted in a yearly mean aboveground biomass accumulation of 5.5 kg/yr, suggesting a mean reduction of 3.5 kg per tree in potential aboveground biomass for the year following the hurricanes. At the stand level, aboveground biomass accumulation across all sites dropped by 4.5 Mg ha^-1^ yr^-1^, from 27.5 Mg ha^-1^ yr^-1^ before the storm to 23 Mg ha^-1^ yr^-1^ after (Figure 5, Table 4). While most forests decreased accumulation rates following the storms, forests in Levittown, Ponce, and of mixed species increased rates. None of the differences, however, between before and after stand level aboveground biomass accumulation were significant.

**Figure 5.**
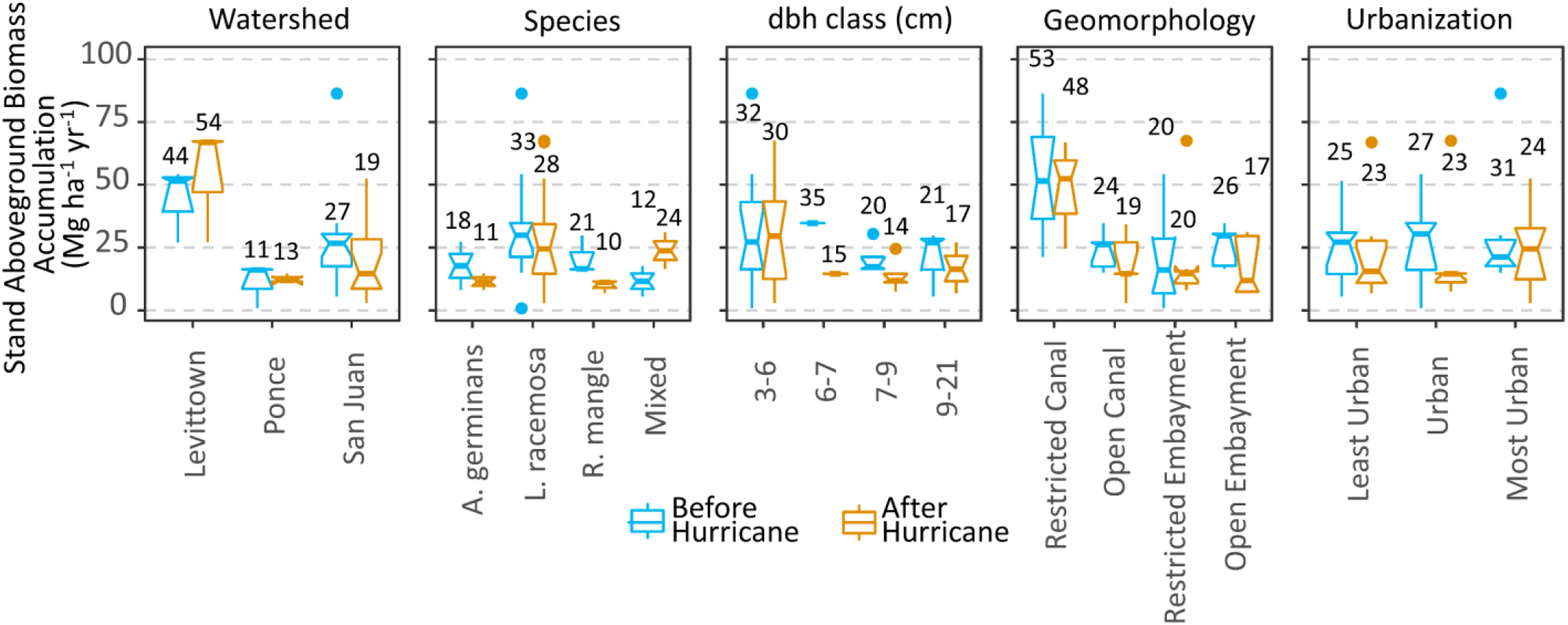
Stand level aboveground biomass accumulation across all groupings before and after the hurricanes. Before values are the means of two measurements taken before the storms. After values are integrations of the curves from Figure 4. Median values are indicated by the bar in the boxplots and means are written above each box. Mean aboveground accumulation dropped across most sites, but increases in Levittown, Ponce, and in mixed forests suggest post-disturbance as outpaced pre-storm forests. None of the differences between pre and post-hurricane aboveground biomass accumulation were significant.

**Table 4.**
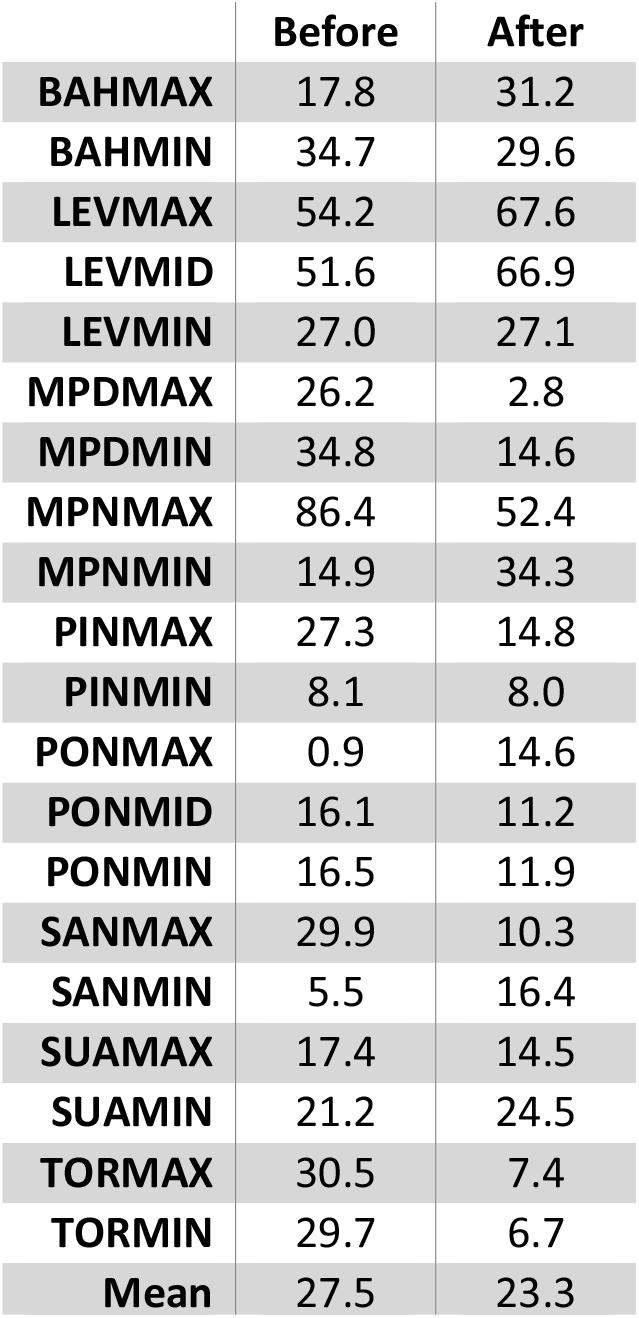
Stand level aboveground biomass accumulation rates before and after hurricanes Irma and Maria in Mg ha^-1^ yr^-1^. In some cases, aboveground biomass accumulation rates have increased, but overall rates have decreased

## Discussion

In the first month following hurricane Maria, the mangroves of Puerto Rico experienced a mean canopy closure loss of 51%, and a mean reduction in aboveground biomass accumulation of 2 kg yr^-1^ per tree (Figure 2). In the following twelve months, 22% of the tagged trees have died, but forest have recovered to 72% canopy closure and to nearly 60% of their pre-storm growth rates. There was only one detected difference between the most urban and least urban sites in the tested metrics, suggesting urbanization plays a minimal role in the tropical cyclone resistance and resilience of Puerto Rico’s mangroves. Instead, tree species, size, and hydro-geomorphic setting were found to explain many of the detected differences between forests. Overall, *L. racemosa* suffered minimal mortality and canopy loss, and recovered more quickly in comparison to the other species. *A. germinans*, however suffered the greatest mortality and canopy loss, and along with *R. mangle*, is recovering slower than the other species. All metrics continue to be monitored and their initial patterns, alongside previous findings, will help determine system recovery over the coming years as well as guide mangrove management to maintain protective services to densely populated areas following tropical storm disturbances.

Differences in initial canopy loss and mortality between species were substantial, and in some cases statistically significant. Mortality results likely underestimate true values due to low sampling size and an inability to account for smaller trees but are still likely indicative of overall patterns. Survival probability remained high until around eight months following hurricane Maria, when it began to drop more quickly. This may reflect inadequacies in the mortality detection method, in which death was only confirmed after all surficial visible signs confirmed it, when in reality trees may have ceased biological function long before (Dobbertin 1998). It may also reflect a lag in tree death following acute disturbance (Filip et al. 2007). In any case, *A. germinans* fared the poorest in this study, and *L. racemosa* the best. The former suffered greater mortality than either *R. mangle* or *L. racemosa*, and stands dominated by it lost about 60% of their canopy closure on average, 13% more than the others. The correlation between canopy closure following the storm and *A. germinans* was strong enough that it could be significantly modeled to decrease by 2% for every 10% increase in the percentage of stand biomass represented by this species. *A. germinans* was also found to suffer the greatest mortality following hurricane Georges in the Dominican Republic (Sherman et al. 2001), but not following hurricane Andrew in Florida (McCoy et al. 1996) or hurricane Hugo in Guadalupe (Daniel Imbert et al. 1996). In parallel, *L. racemosa* has been shown to be both the most resistant (Armentano et al. 1995; Sherman et al. 2001), as well as the most susceptible species to hurricane mortality (Wadsworth 1959; Smith et al. 1994; McCoy et al. 1996). These contradictions may come from differences in how mortality was determined and after differing lengths of time, but also from differences in habitat types, which has been found to significantly influence interspecific mortality (Armentano et al. 1995; Smith III et al. 2009). This might explain why open embayment systems suffered less canopy loss and mortality than tidally restricted canal systems in this study.

As with species, size was another important explanatory variable in initial mortality. Larger individuals of all species suffered greater mortality than smaller individuals. A number of studies have shown that large mangrove trees (dbh > 10cm) are more susceptible than smaller trees to canopy loss and mortality following hurricanes (Roth 1992; Smith et al. 1994; McCoy et al. 1996). The consistency of this pattern among previous studies as well as this one, gives further weight to the hypothesis that Caribbean mangrove height is partly dependent upon hurricane frequency, with larger trees selected against due to their greater susceptibility (Odum and Pigeon 1970; Lugo and Snedaker 1974; Doyle and Girod 1997). Thus, it seems pertinent to consider larger trees at a greater risk to hurricane mortality, and this should be considered when evaluating the potential loss of mangrove ecosystem services along densely populated shorelines.

With a mean mortality of 22%, Puerto Rico’s mangrove’s seem to have fared better than those after other storms, whose mortality ranged from 25% to 90% (Craighead and Gilbert 1962; Roth 1992; Smith et al. 1994; Armentano et al. 1995; Sherman et al. 2001; Daniel Imbert 2018). Although this point may be due to differences in survey methodologies, the definition of “mortality”, and/or study lengths between studies, Puerto Rico’s presence at the far lower extreme of this range is notable. While partial and complete mortality following the hurricane is likely due mostly to interspecific and inter-size differences in susceptibility, as well as some contribution from geomorphology, it does not seem to be influenced by urbanization. Distance or wind energy also were not significant predictors of tree death, canopy loss, or recovery. This is surprising given the differences in wind power between Ponce and the northern coast sites (Van Beusekom et al. 2018), which is consistent with a lower mortality at Ponce. This may reflect an inadequate tree sampling size and/or inaccuracy in the wind power model.

As with initial mortality and canopy loss, size and species were significant predictors of differences in mangrove recovery across the forests. Non-mangrove species grew faster than mangroves (Figure 4). This may be explained by the extended depth and presence of freshwater lenses in the mangroves following the storms. This freshwater, along with an excess of understory sunlight, allowed existing non-halophytes to thrive (Lugo 1999). As for mangroves, *L. racemosa* grew faster than both *R. mangle* and *A. germinans*. *R. mangle’s* failure to grow is likely explained by its diminished epicormic re-sprouting abilities and its ground-up regeneration strategy (Wadsworth 1959; Tomlinson 1980; A. Baldwin et al. 2001). *Avicennia spp*., however, have repeatedly been found to be of the most resistant species to hurricane disturbance (Woodroffe and Grime 1999; Daniel Imbert 2018), so its failure to regrow in this study is contradictory. It’s possible that because all sites in this study were fringe systems of low *A. germinans* density (Branoff and Martinuzzi 2018), recovery following the storm was made more difficult by stressful and unsuitable habitat. Patterns in size class growth rates suggest that although the largest trees continued to accumulate more aboveground biomass than smaller trees following the storm, their growth rates steadily diminished with time (Figure 4). Smaller trees, however, were accumulating far more in respect to their own biomass, suggesting recruits are taking advantage of excess sunlight and have begun competing for canopy space (A. Baldwin et al. 2001; Ward et al. 2006; Daniel Imbert 2018).

Hydro-geomorphology was also a consistently significant predictor of differences in initial mangrove mortality and subsequent recovery. Initial canopy loss was greatest in restricted systems with only partial tidal connectivity (Figure 2), but trees in these forests then grew quicker than those in other forests following the storms (Figure 4). This may be due to differences in hydrology and surface water chemistry in these forests, both of which are known to play important roles in mangrove function (Lugo and Snedaker 1974; Wolanski et al. 1993). Other studies have shown riverine and fringe mangroves to suffer less mortality than basin systems (Smith III et al. 2009; Daniel Imbert 2018), which is likely due to quicker drainage time following storm surges, and thus lower hypoxia related stress to roots. The tidally restricted systems of this study may also share this benefit as storm surges may not have reached as far as more tidally open systems.

As the forests continue to recover one year after hurricane Maria, canopy closure will likely be one of the most important determining factors in successional and structural dynamics (Muscolo et al. 2014). In this study across all forests, closure to 80, 90, and 95 % could be predicted to occur within 3.6, 9.7, and 16 years, respectively. This agrees with the 8-14 years predicted for gap closure across multiple forest types (Runkle 1981; Horvitz and Schemske 1986; Cipollini et al. 1993; Valverde and Silvertown 1997). Closure in forests dominated by *A. germinans* and *R. mangle*, however, could not be reliably forecasted because of their high mortality and low regeneration rates. Instead of canopy closure through existing tree growth in these forests, gaps will likely experience high recruitment rates that will result in a longer canopy closure timeline (Lugo et al. 1976). In the meantime, although most forests have experienced a dip in stand level aboveground biomass accumulation following the storms (Figure 5), some have increased accumulation and will likely continue to do so as the post-disturbance environment favors high recruitment and growth in surviving trees. As a result, post-hurricane forests may not resemble their pre-storm characterizations, and will likely instead experience shifts in species distributions and structure that persist for decades if not centuries (Smith et al. 1994; A. Baldwin et al. 2001; Daniel Imbert 2018).

Although urbanization has been found to be influential in forest ecology and disturbance, this study found little evidence of such an influence. Highly urban mangrove forests could be deemed neither more nor less susceptible to hurricane mortality or canopy loss. Instead, the usual suspects of species, size, and geomorphology, were more strongly identified as influential in determining initial response and short-term recovery of hurricane disturbed mangroves in Puerto Rico. This implies that it may not be necessary to strongly consider surrounding urban land cover in the management of mangroves for optimal protective services. When shoreline protection and stabilization is by far the most valuable service provided by mangroves (Costanza et al. 2008; de Groot et al. 2012), and in urban settings where there is more life and property to protect, optimizing this service may simply mean managing forests to promote smaller individuals of *L. racemosa* in restricted canal geomorphologies. But the above stated inconsistencies between the findings of this study and those of others, point to a need for more studies of mangrove ecology along well-defined urban gradients. Such studies include the continued monitoring of these forests for long-term successional and recovery dynamics, as well as pre-storm baseline measurements in strategic locations within tropical cyclone prone areas. Doing so will provide much needed information on the role of social influences, in addition to ecological ones, in the protective services of mangroves, thus allowing managers to make more informed decisions towards optimizing social-ecological mangrove ecosystems.

## ACKNOWLEDGEMENTS

Research was partly funded by the Next Generation Ecosystem Experiments-Tropics, funded by the U.S. Department of Energy, Office of Science, Office of Biological and Environmental Research and by. NASA’s Goddard Space Flight Center coordinated LiDAR flights. Ariel Lugo of the US Forest Service International Institute of Tropical Forestry provided feedback on first drafts.

